# trAIt: Species-by-Trait Data Retrieval using Large Language Models

**DOI:** 10.64898/2026.06.19.732660

**Authors:** Srivi Balaji, Katherine A. Martinson, Jessica S. Schellenberger, Jyotishko Koley, Callen M. Inman, Hans A. Hofmann, Rebecca L. Young, Arbel Harpak

## Abstract

Biological research often requires information about species’ traits. Manual literature collation can be time-consuming and miss parts of the literature. To address this gap, we developed ***trAIt***, a publicly available software for the retrieval of characteristics of species from scientific literature catalogued in the Europe PubMed Central (PubMed) database. *trAIt* provides a graphical user interface (GUI) in which users specify species and characteristics of interest. Leveraging a large language model (LLM), *trAIt* retrieves relevant papers, combines their content through a consensus-based summarization model, and outputs a species-by-characteristic table. For a case study involving frog species, *trAIt* recovered 47.1% of trait-species combinations in 2.75 hours, while an expert curator independently recovered 62.4% over months. The consensus-based summarization substantially aids accuracy compared to single-source extraction. Across three case studies of vertebrate taxa, an expert confirmed the accuracy of 70.9% of trait-species entries recovered by *trAIt*. We observed considerable variation across taxa in *trAIt*’s accuracy, which is possibly due to heterogeneity in open-access literature availability and inconsistencies in species and trait terminology. In sum, our analysis suggests that LLM-based tools can accelerate biological data synthesis but should be used to support domain experts’ research, rather than replace their judgment.

## 1 Introduction

Data on species, such as their morphological, behavioral, and ecological characteristics, is central to many areas of biological research. These data may be derived from expert observations, field studies, or laboratory experiments. They can be acquired from several sources, including public databases, primary literature, and code repositories, to name a few. Yet researchers face obstacles during data collection. For example, in human studies, both interviewer and interviewee can experience fatigue when conducting interviews or surveys^1^. Additionally, variation in formatting or annotation practices can make some sources difficult to interpret and compare^2,3^. These challenges present the opportunity to streamline literature-based data retrieval.

One approach for faster trait data retrieval is to use artificial intelligence (AI) models. We developed *trAIt*, a trait-by-species data retrieval software. *trAIt* queries scientific literature and leverages a large language model (LLM) to extract concise information from large bodies of text. It then summarizes answers across sources and outputs a trait-by-species table. In designing and evaluating *trAIt*, we paid special attention to “old” problems experienced by human researchers and “new” problems we expected LLM chatbots might introduce. As an example of an “old” problem, we considered inconsistent trait nomenclature. In frogs, morphological size may be referred to as “Body Size,” “Snout-Vent Length,” or “Adult Length”. This impedes manual synthesis because it requires identifying and searching for multiple naming variants to capture all available information. It was not clear to us at the outset whether this would pose a bigger problem to human experts or chatbots. As an example of a new problem, AI chatbots are fast but not necessarily accurate or reliable. To make matters worse, chatbots are infamous for sometimes being deceptively confident about answers they “hallucinate”^4^. To be an effective tool for researchers, *trAIt* ought to address these challenges.

Here, we describe *trAIt*’s design aimed at hitting the aforementioned goalposts. We evaluate performance on multiple test cases in terms of speed, completeness, and accuracy. Lastly, we ask whether *trAIt* could be improved by using a taxon-specific database for data extraction rather than primary literature. We explore the potential benefits of using *trAIt* and discuss the limitations of species-by-character data extraction using *trAIt*, and more generally, pitfalls and windfalls of AI use in literature surveys.

## 2 Results and Methods

### 2.1 System Overview

*trAIt* uses a workflow (**Fig. 1**) with the following steps. See **Table 1** for definitions.

**Figure 1.**
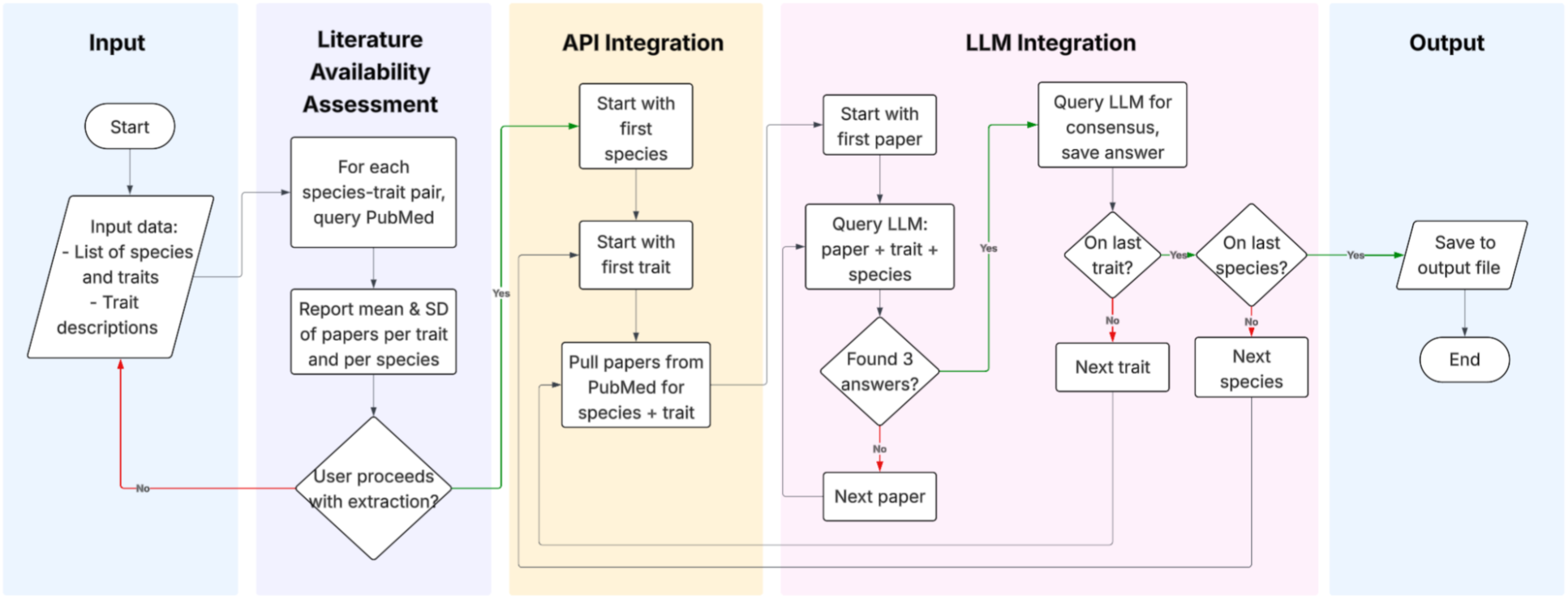
Overview of the generalized species-trait retrieval pipeline. For each species-trait pair, *trAIt* queries PubMed, extracts relevant text, and uses the configured LLM to identify and summarize trait information.

**Table 1.**
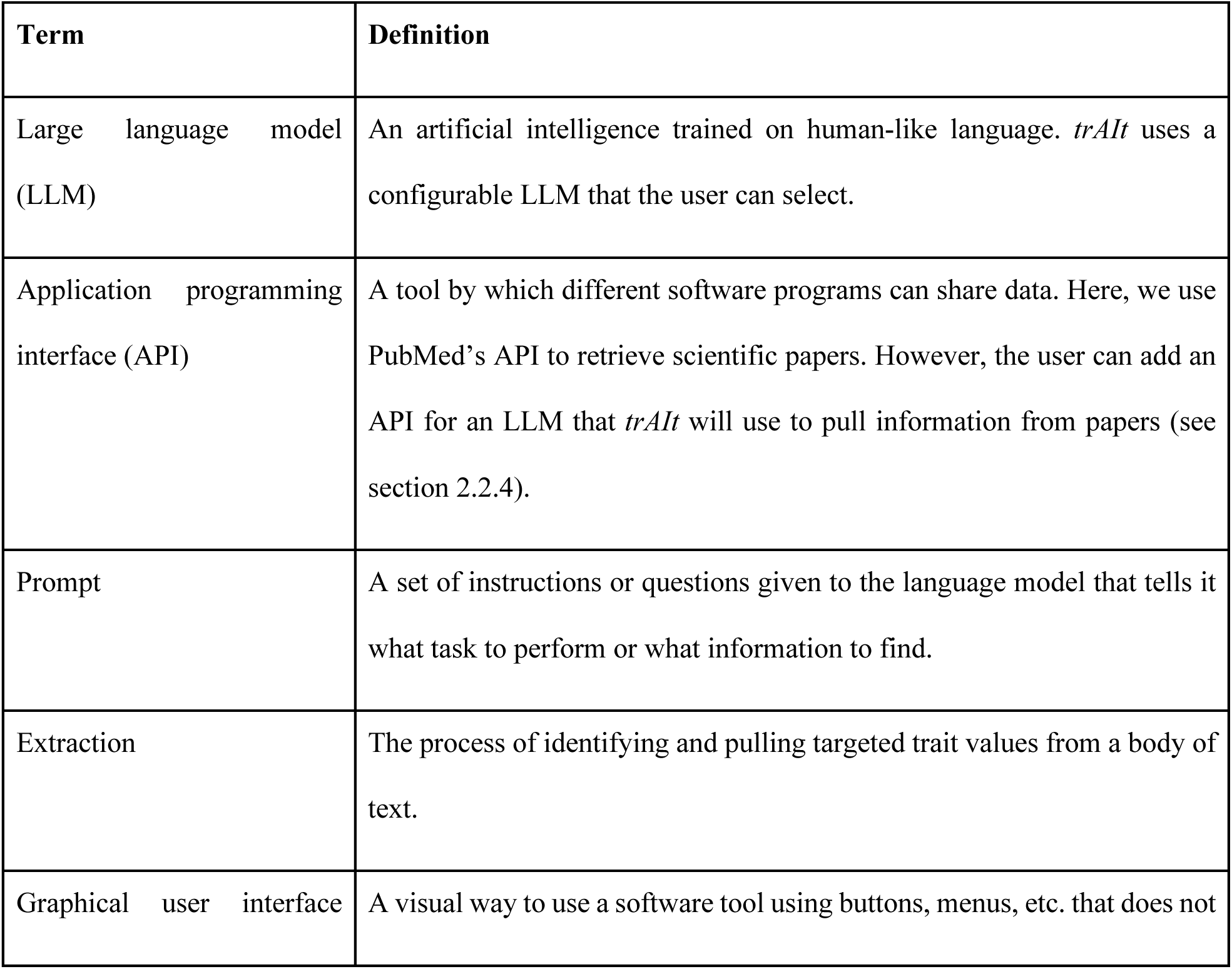
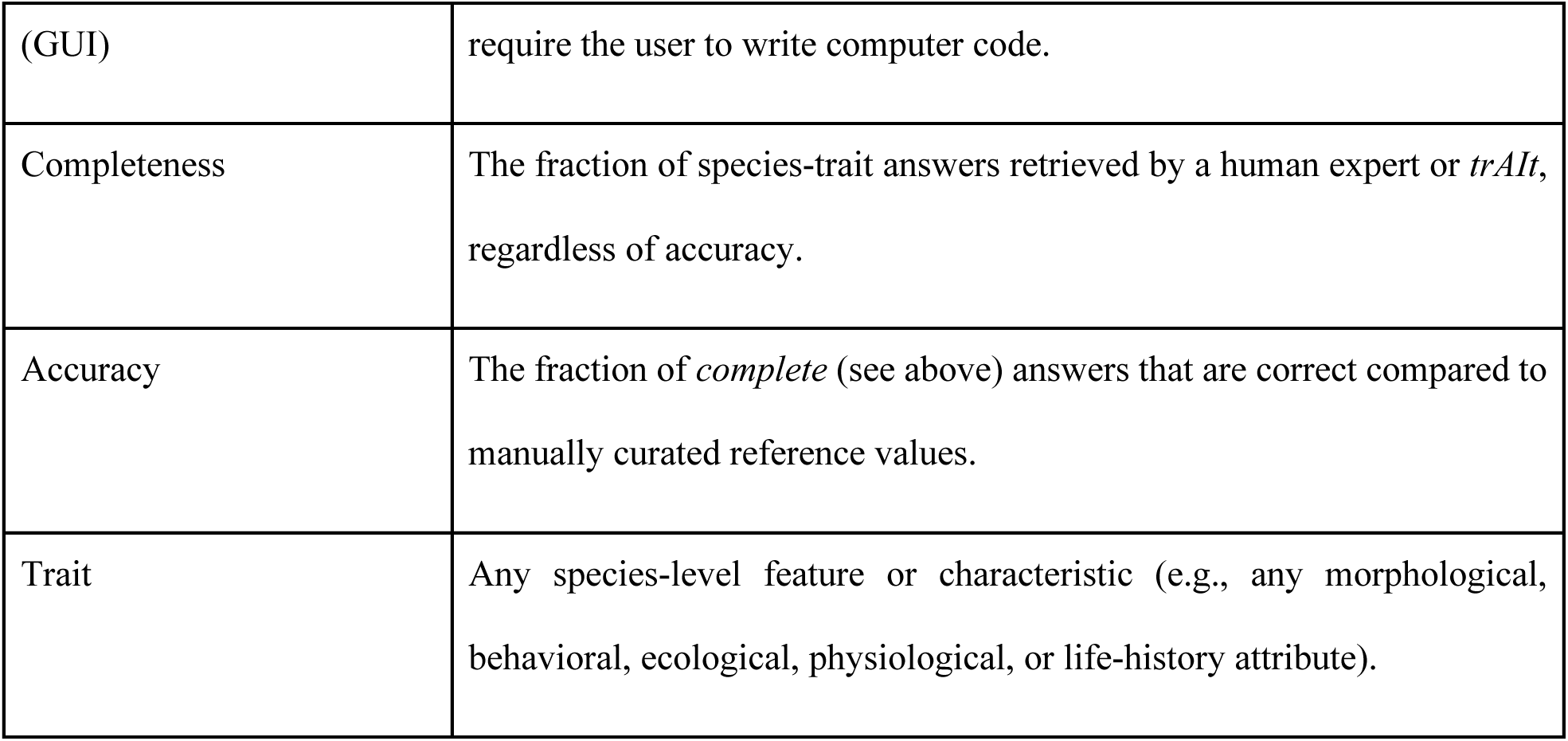
Key terms and definitions.

1. **User Input:** The user uploads a spreadsheet listing species names and traits of interest, along with a text file describing each trait in plain language.
2. **Literature Retrieval:** For each species-trait pair, *trAIt* retrieves relevant papers for that species and trait through PubMed’s open-access API.
3. **Text Processing:** *trAIt* downloads retrieved papers in PDF format and converts them to raw text. It ensures data quality by filtering incomplete or unparseable documents and preparing clean text for analysis.
4. **Iterative LLM Querying:** For each retrieved paper, *trAIt* applies a standardized extraction prompt to the LLM, supplying the paper text and trait definition, and receiving the model’s response corresponding to the requested trait. Before running *trAIt*, the user can specify the LLM they want to use by adding an API key to a configuration file.
5. **Consensus Summarization:** If a query returns multiple candidate values from multiple papers, *trAIt* reconciles and standardizes the results into a single, unified output.
6. **Results Compilation:** The aggregated results are written to a structured CSV file, and logs record all papers reviewed and those that yielded relevant trait information.

### 2.2 System Components

#### 2.2.1 Input

The user provides two inputs:

- A spreadsheet containing species names in the first column and requested traits in subsequent columns.
- A text file that defines each trait in simple language.

The GUI (**Fig. 2A**) guides the user through preparation of these files, offering example templates and reference images to ensure correct formatting. After inputs are uploaded, the interface displays progress messages and directs the user to the final output files once processing is complete.

**Figure 2.**
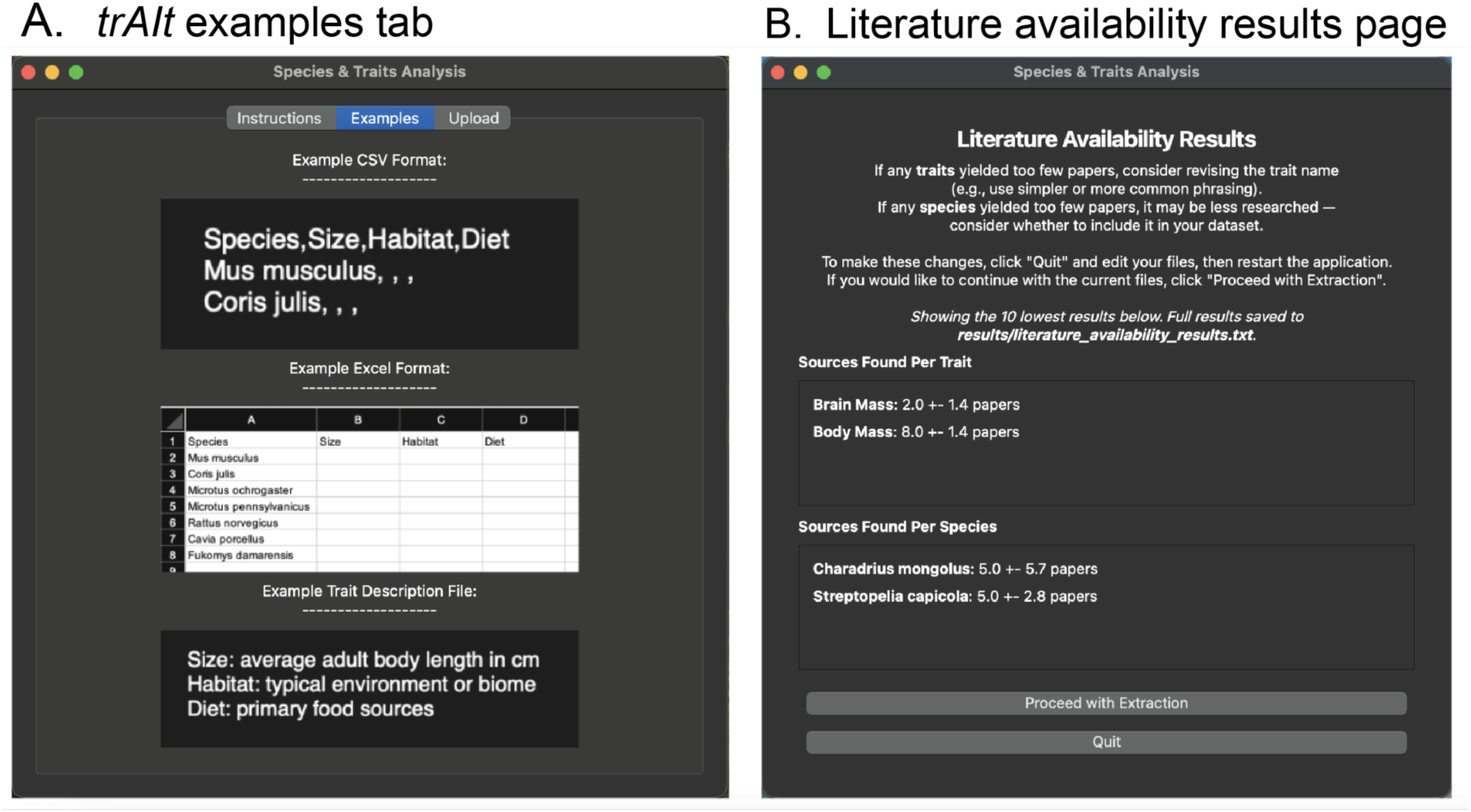
GUI for *trAIt* input preparation and literature availability assessment. **(A)** The interface provides sample CSV, Excel, and trait-description formats to illustrate how the user should structure their data prior to running *trAIt*. **(B)** After inputs are uploaded, *trAIt* displays literature availability metrics for each trait, showing the mean number of papers retrieved and the standard deviation across all queried species and traits.

#### 2.2.2 Literature Availability Assessment

Before beginning the extraction step, *trAIt* assesses the availability of literature for each requested species and trait (**Fig. 2B**). This gives the user the opportunity to evaluate whether the amount of available data justifies proceeding, and to troubleshoot species names, trait names, or trait descriptions that yielded few results. For each trait and species in the user’s input file, *trAIt* queries PubMed and reports the mean and standard deviation of the number of papers across species and traits individually^5^. The user can then choose whether to proceed or revisit and modify their input. This evaluation helps avoid unnecessary computational costs and runtime which can be substantial in the LLM integration step.

#### 2.2.3 Literature Retrieval and Text Processing

*trAIt* retrieves relevant scientific literature for each species-trait pair through the PubMed open-access API. This stage includes several steps:

1. **Query Construction:** For each species-trait pair, *trAIt* formulates a structured search query designed to retrieve papers in which the terms co-occur in biologically relevant contexts.
2. **Source Retrieval:** *trAIt* searches the PubMed database, which aggregates open-access research articles across biomedical and ecological disciplines.
3. **Paper Selection and Filtering:** *trAIt* filters retrieved papers to ensure relevance and completeness. It excludes papers that cannot be downloaded, lack full text, or produce parsing errors (e.g., incorrect file formats, PDF corruption or encryption, or empty text extraction).
4. **Text Preparation:** *trAIt* downloads and converts full-text articles (PDF or XML) into plain text by stripping formatting, figures, and layout elements to produce a clean text block for LLM analysis. *trAIt* also manages text length constraints, ensuring that documents exceeding predefined token limits (i.e., documents that are too long for the LLM to read) are clipped while retaining important information.

#### 2.2.4 LLM-Based Trait Extraction

After preprocessing, *trAIt* passes the cleaned text to the LLM. For each species-trait pair, the LLM receives as input:

- the extracted text from the paper, and
- a clear natural-language instruction based on the trait definition provided by the user.

#### 2.2.5 Consensus Model

When *trAIt* queries the literature for a single species-trait pair, it retrieves up to twenty candidate papers. The LLM attempts to extract trait values from each paper. The search is completed when *trAIt* collects an answer from a total of three separate papers or when the model reaches twenty total papers.

For the consensus model, *trAIt* combines up to three answers using a consensus mechanism:

- When values are similar (e.g., 10 cm, 11 cm, and 10.5 cm), it produces a range (“∼10-11 cm”).
- When multiple categorical answers are retrieved (e.g., “temperate forests, wetlands”), it merges them into a composite answer separated by commas.
- It discards outputs that fail basic validation (e.g., illegible text) to avoid polluting the output.

If only one retrieved paper provides a validated trait value, *trAIt* records that single value without further aggregation. If no papers provide information, *trAIt* marks the species-trait entry as missing in the final output.

#### 2.2.6 Output Compilation

After consensus resolution, *trAIt* writes the aggregated trait values for all species into a structured CSV file. *trAIt* outputs a log file that records *all* papers that were retrieved when predicting each species-trait entry, and a success log file that records *only* papers where the LLM returned a non-missing trait value for that species-trait entry. The success log contains a maximum of three papers per species-trait pair, sometimes fewer when literature availability is low.

### 2.3 Performance Evaluation

To assess the performance and generality of *trAIt*, we applied it to three inputs (**Table 2**): a *Mixed Vertebrate* input consisting of 23 vertebrate species and 14 traits; a *Frog* input consisting of 200 species and 12 traits; and a *Bird* input consisting of 146 species and 9 traits. These inputs were filled in manually by experts in each of their respective clades.

**Table 2.** Summary of evaluation data. SVL: Snout-Vent Length.

| Input | Number of Species | Traits |
| --- | --- | --- |
| Mixed Vertebrate | 23 | Mating System, Social Dominance Hierarchy, Territoriality (Males), Territoriality (Females), Group Size During Reproduction, Group Size Outside of Reproduction, Group Property, Age at Maturity, Average Life Expectancy, # Offspring/Reproductive Bout, Reproductive Rate, Migratory Behavior, Activity Pattern, Habitat Complexity |
| Frog | 200 | SVL Male, SVL Female, Average SVL Adult, Egg Clutch, Average Temperature, Average Rainfall, Average Altitude, Egg Style, Average Hatch Time, Average Development Time, Average Time of Day, Average Age at Maturity |
| Bird | 146 | Body Mass, Brain Mass, Colonial, Maximum Colony Size, Cooperative Breeding, Communal Foraging, Sociality Outside Of Breeding, Migratory, Habitat Type |

**Table 3** shows an example of the output format of *trAIt* for a subset of the Mixed Vertebrate input. The trait descriptions provided to *trAIt* for these example traits were as follows:

**Table 3.** Example *trAIt* output spreadsheet.

| Species | Mating System | Social Dominance Hierarchy | Territoriality (males) | Territoriality (females) | Group Size During Reproduction |
| --- | --- | --- | --- | --- | --- |
| *Microtus ochrogaster* | Monogamous | Linear | Small territories | Small territories | 2-10 individuals |
| *Microtus pennsylvanicus* | Promiscuous | No | Small territories | Small territories | solitary |
| *Mus musculus* | Monogamous | Linear | Variable territory size: small to large | No territoriality observed. | 2-10 individuals |
| *Rattus norvegicus* | Polygynous, Monogamous | Non-linear |  |  | 2-10 individuals |
| *Cavia porcellus* | Promiscuous, Monogamous | Linear | Large territories |  | 2-10, 10-100 |
| *Fukomys damarensis* | Monogamous | Linear | No |  | 2-100 individuals |
| *Pan troglodytes* | Promiscuous | Non-linear, Linear | Large territories | Small territories, No territoriality |  |
| *Homo sapiens* | Promiscuous, Monogamous | Non-Linear, Linear |  | Large territories, no territoriality (females disperse) | 100-1000+ individuals |
| *Gorilla gorilla* | Polygynous | Linear | Small territories, No | No, Small territories | N/A |

- **Mating System:** The structure of social and sexual interactions of reproduction. This is a categorical trait, and the options are “Monogamous”, “Polygynous”, “Polyandrous”, “Polygamous”, and “Promiscuous”.
- **Social Dominance Hierarchy:** How non-mating individuals in a group interact with one another. This is a categorical trait, and the options are “No”, “Linear”, and “Non-Linear”.
- **Territoriality (males):** Whether males of this species maintain territory, and how much territory. This is a categorical trait, and the options are “No”, “Small territories”, and “Large territories”.
- **Territoriality (females):** Whether females of this species maintain territory, and how much territory. This is a categorical trait, and the options are “No”, “Small territories”, and “Large territories”.
- **Group Size During Reproduction:** The size of the social group while reproducing. This is a categorical trait, and the options are “Solitary (no sociality)”, “Small groups (2-10 individuals)”, “Medium groups (10-100 individuals)”, and “Large groups (100-1000 or more individuals)”.

All five of these traits are categorical, meaning *trAIt* extracts one of a predefined set of category labels for each species. For example, Group Size During Reproduction has four possible categories: “solitary”, “2-10 individuals”, “10-100 individuals”, and “100-1000+ individuals.” Note that for categorical traits, we used trait descriptions that explicitly list the values we expected. For all quantitative and qualitative evaluations reported in this section, *trAIt* was executed using GPT-5-nano as the configured LLM^6^.

#### 2.3.1 Comparison of Extraction Strategies: First-Hit vs Consensus

We first compared two model architectures for trait extraction: a **first-hit model**, which records the first extracted value encountered across retrieved papers, and a **consensus model**, which aggregates outputs from multiple sources and combines them into one trait answer. Papers are retrieved in order of relevance, which is a score given to each document that factors in the frequency of the search terms within the document, how rare the terms are across all documents, and how recent the document is^5^.

In the Mixed Vertebrate input, the consensus model consistently produced more accurate and complete outputs. The first-hit method often captured partial or context-specific values like a measurement reported for one study site or population, whereas the consensus approach reconciled conflicting measurements and incorporated multiple study contexts. An example of this was when *trAIt* attempted to fill in data for model organisms, such as *Mus musculus* and *Homo sapiens*. The first-hit model would often grab results from clinical studies that do not represent the intended wild population data. The consensus model significantly improved average accuracy from 50.0% to 75.2% (**Fig. 3**, p-value < 1×10^-4^, paired t-test). Because the consensus approach consistently outperformed the first-hit strategy in the Mixed Vertebrate input, we conducted all subsequent evaluations only using the consensus model.

**Figure 3.**
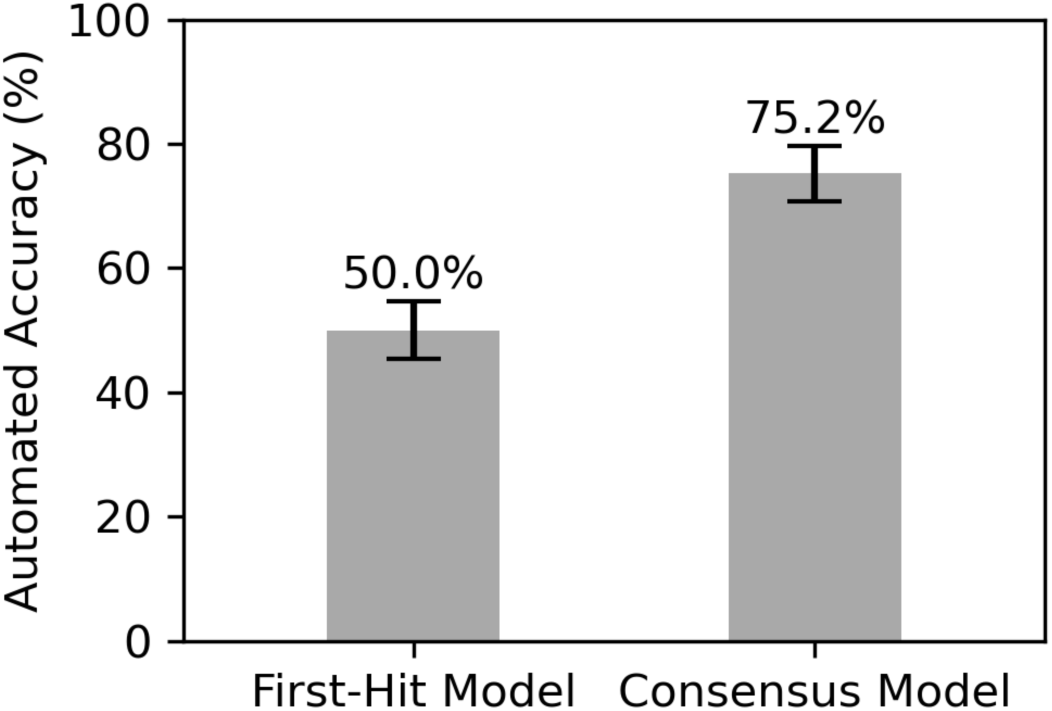
Accuracy comparison between two *trAIt* models on the Mixed Vertebrate input. In the first-hit model, 50% of the retrieved entries were accurate. In the consensus model, 75.2% of the retrieved entries were accurate. Automated accuracy is estimated using the proportion of non-missing trait-by-species outputs that match manually curated reference values (via exact text match). Bars represent +/- SE across species. The consensus model significantly outperforms the first-hit approach.

#### 2.3.2 Qualitative Evaluation

We found *trAIt* to be robust to nomenclature heterogeneity that makes manual literature synthesis challenging. *trAIt* recognized that trait names “Snout-Vent Length”, “Adult Body Length” and “SVL” are used synonymously and extracted information accordingly. At the same time, there were instances in which it was misled by acronyms or abbreviations that would not have misled a human expert. For example, *trAIt* treated “Average Life Expectancy” and the abbreviated “Avg. Life Expectancy” as distinct phrases, retrieving papers for the former but missing the same papers for the latter.

Similarly, semantic ambiguity remained a challenge. When extracting “Colonial Nesting” as a trait for the Bird input, we noticed that papers sometimes used broader descriptors, like “congregatory” or “social” nesting, which are not specific enough to determine whether the species nested colonially. Yet *trAIt* might determine that these terms are semantically similar enough and extract an answer. This can lead to extractions that may be correct but are too general for the intended trait definition. On the other hand, *trAIt* can also miss correct information due to incorrect semantic interpretation. For example, when querying “cone opening ability” (the ability of rodents to forage for pinecones) from a rodent input not included in this analysis, *trAIt* retrieved many papers in the literature availability assessment. However, the LLM interpreted “cone” in the visual sense, looking for information about cone opsins instead of foraging behavior. This demonstrates that high paper counts in the literature availability assessment do not guarantee that the trait name will be interpreted correctly by the LLM.

A key limitation is that trait values are often reported in figures or supplementary materials rather than in the source text itself. Because *trAIt* extracts only text from the main body of papers, it may return missing values for species-trait pairs where the relevant information exists in the paper but not in the main text. We also observed challenges with output format consistency for categorical traits. For example, when extracting “Group Size During Reproduction”, the LLM sometimes returned “2-10” and other times “2-10 individuals” for the same category. While it is difficult to tune the LLM to always produce the same format, a user can address this through post-processing steps to standardize category values.

#### 2.3.3 Quantitative Evaluation

We considered both the completeness and accuracy of information retrieved by *trAIt*. Completeness varied considerably across inputs—50.6%, 60.1%, and 26.6% for Mixed Vertebrates, Birds and Frogs, respectively (**Fig. 4A**).

**Figure 4.**
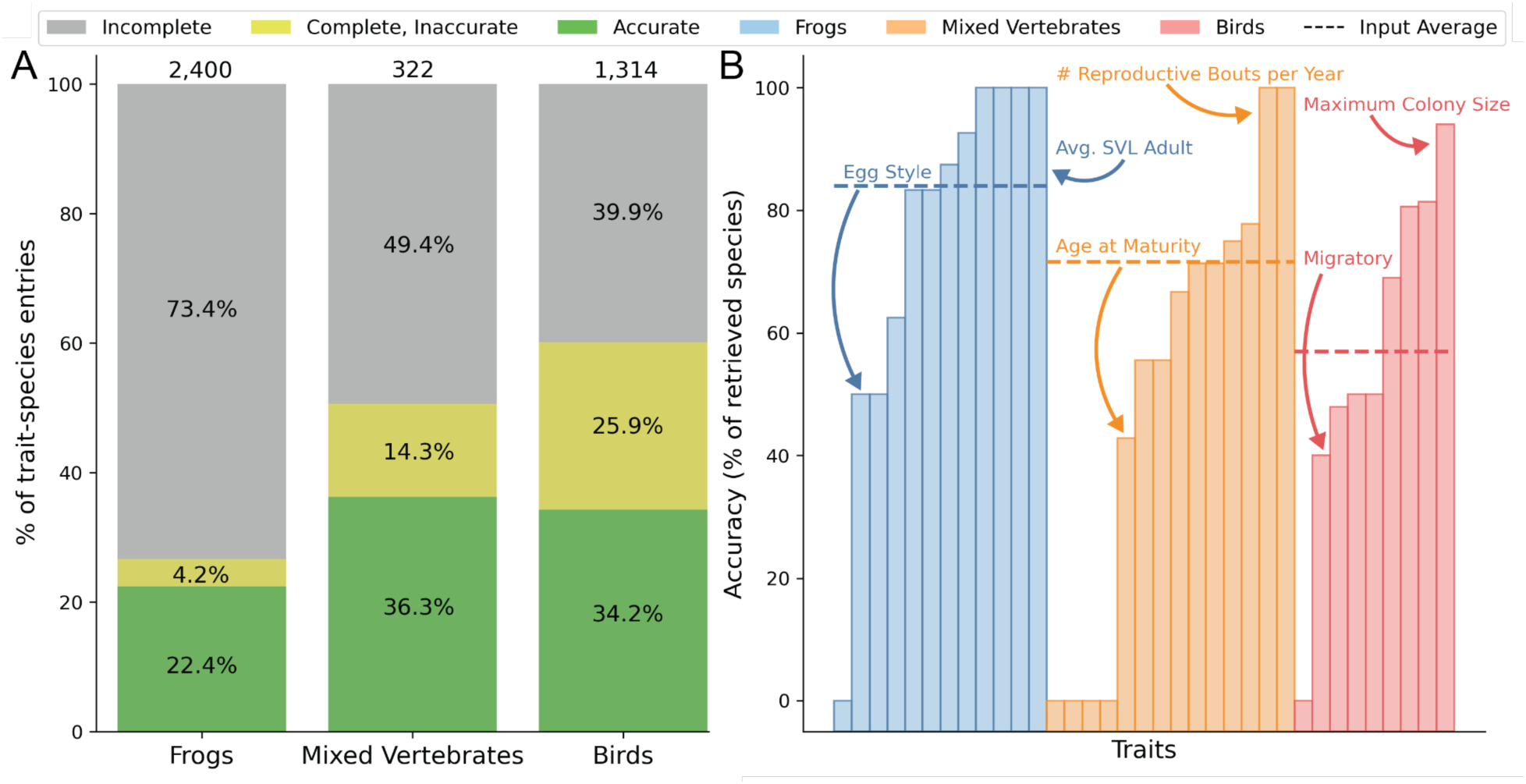
Quantitative evaluation of *trAIt’s* performance. **(A)** Average accuracy and completeness across all trait-species entries across traits. Percentages within the bars represent the proportion of trait-species entries that were returned by *trAIt* as incomplete (gray), complete but inaccurate (yellow), and accurate (green). Numbers on top of each bar represent the total number of trait-species pairs. **(B)** Distribution of accuracy, evaluated by a human domain expert, across traits. See **Fig. S1** for analogous results when using automatic (computerized) evaluation of accuracy. Input average is the average accuracy across all traits across an input.

To evaluate the accuracy of *trAIt*’s output, we compared it to a trait-by-species table manually compiled and curated by the authors (as domain experts) before running *trAIt*. We only considered species-trait pairs for which *trAIt* produced an answer (“complete”). For categorical traits, we assess accuracy by a direct text match. For quantitative traits, we consider *trAIt*’s answer as accurate if it is less than 1.5 standard deviations away from the consensus output, calculated across all species in the ground-truth input. For traits where *trAIt* extracted a range, we considered the output correct if the reference value fell within that range.

Accuracy showed similar distributions across traits within each input (**Fig. 4B**). Some of the most accurate traits across all inputs are those involved with reproduction or behavior (Brain Mass, Colony Size, # Reproductive Bouts Per Year).

Among the three inputs, *trAIt* achieved an accuracy of 71.6% on the Mixed Vertebrate input, 57.0% on the Bird input, and 84.0% on the Frog input (**Fig. 4A**). These results highlight that *trAIt*’s performance is not uniform across taxa. Accuracy depended on the species and traits being extracted, likely reflecting variation in literature availability and reporting standardization across taxonomic groups^7^. Several species in the Mixed Vertebrate input, like *Mus musculus* and *Rattus norvegicus*, are extensively studied, with multiple well-annotated papers that report trait values in clear, accessible formats^8^. This is a plausible reason for the higher automatic text-match accuracy of *trAIt* in this case (74.1%, **Fig. S1A**). However, because some vertebrates also appear frequently in clinical or laboratory studies, retrieved papers may describe traits measured under experimental or clinical conditions rather than in the wild, which can result in *trAIt* extracting information that does not represent the species’ natural characteristics. Such outputs would be considered inaccurate. The Bird input generally benefited from higher availability of relevant literature compared to amphibians. Per a previous report, there are 345 million bird occurrences in the Global Biodiversity Information Facility (GBIF) database, while amphibians have only 3.94 million occurrences^7^.

### 2.4 Case Study: Taxon-Specific Data Sources

For Frogs, *trAIt’s* output was accurate (83.9%) but highly incomplete (only 26.6% of 2,400 entries; **Fig. 4A**). We hypothesized that this is a result of the limited availability of open-access papers on many of the species in the Frog input. To test this hypothesis, we evaluated completeness when using a more specific source of data. In particular, we replaced the querying of PubMed with the API of AmphibiaWeb, a structured, curated, amphibians-specific database^9^.

AmphibiaWeb provides structured data in XML format for amphibians, while International Union for Conservation of Nature (IUCN) and the Climate Change Knowledge Portal (CCKP) supply environmental, range, and conservation metadata^10,11^. Instead of pulling papers’ text from the PubMed API (**Fig. 1**, “API Integration” box, step 3), *trAIt* retrieved trait-relevant content directly from structured databases: AmphibiaWeb, IUCN, and CCKP. With structured data, we hypothesized that there is a higher likelihood of the data containing the traits we are looking for. Primary literature queries rely on PubMed’s relevance metric, whereas APIs can more directly point *trAIt* to specific sections of the data.

We evaluated *trAIt*’s extraction performance against the same manually curated reference for the Frog input that was used to evaluate *trAIt*’s performance in section 2.3. Completeness rose from 26.6% using PubMed to 47.1% using AmphibiaWeb. Accuracy rose from 84% using PubMed to 89% using AmphibiaWeb (**Fig. 5A,B**). However, this accuracy result may be due to the Frog ground truth, which comes from inputs manually curated by domain experts, being heavily dependent on data from AmphibiaWeb. Given that *trAIt* and the human expert had access to the same data, there is a bias in our accuracy measurement when using AmphibiaWeb.

**Figure 5.**
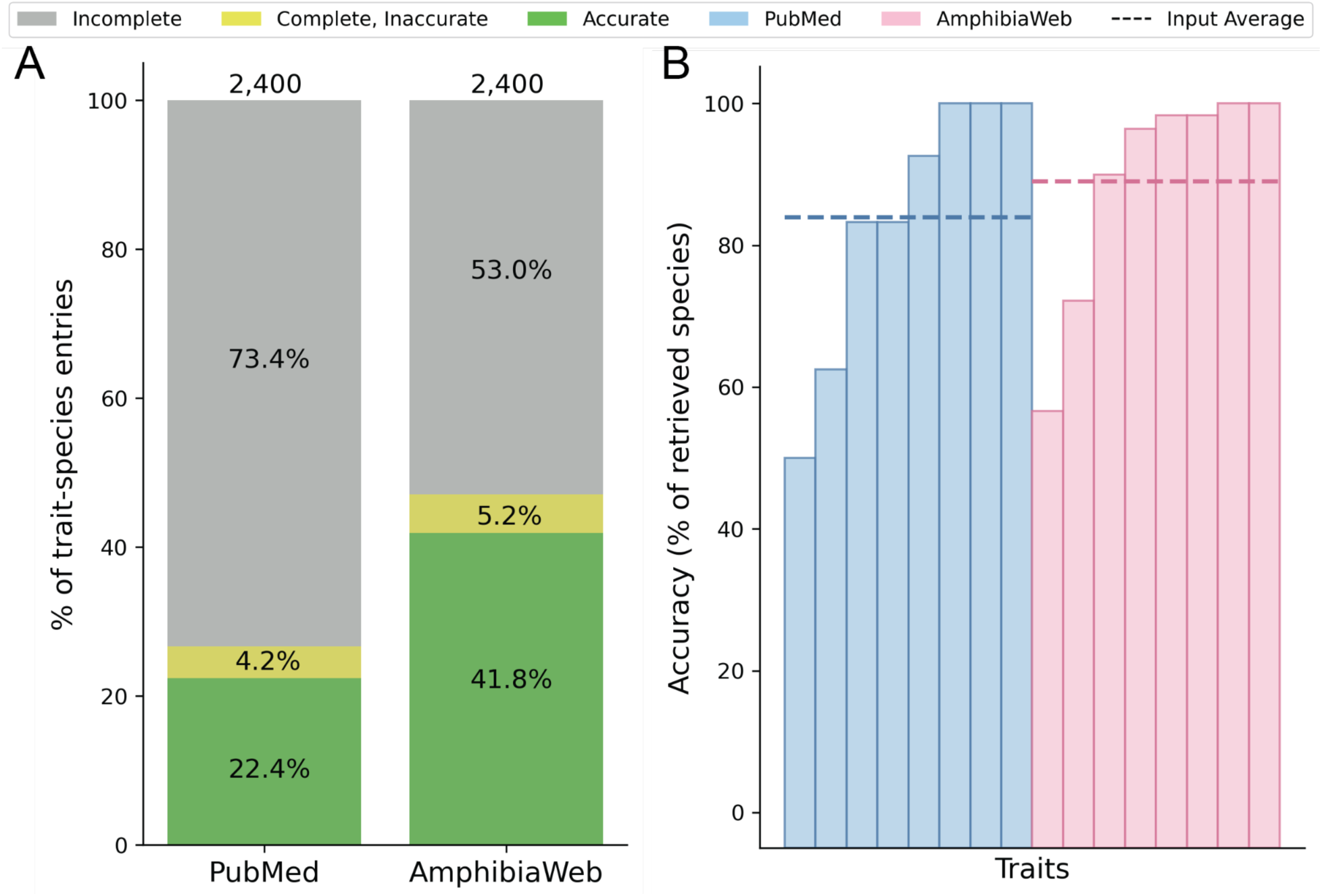
Comparison of *trAIt* extraction performance on the Frog input when using PubMed versus AmphibiaWeb. **(A)** Average accuracy and completeness across all trait-species entries across traits. Percentages within the bars represent the proportion of trait-species entries that were returned by *trAIt* as incomplete (gray), complete but inaccurate (yellow), and accurate (green). Numbers on top of each bar represent the total number of trait-species pairs. **(B)** Distribution of accuracy across traits. Input average is the average accuracy across all traits within sample inputs.

## 3 Discussion

*trAIt* provides an automated approach for extracting biological trait information from scientific literature. We found that *trAIt*’s performance depended heavily on literature availability and reporting consistency. In addition, our analysis of the change in performance when using a taxon-specific database demonstrated that when literature is available in structured text form, *trAIt* can more reliably extract information (**Fig. 5**). Notably, *trAIt* relies on data being mineable as text rather than in figures, files or other supplementary materials. This poses a significant limitation, as it is common practice to report trait values in these forms. While we hope this limitation could be tackled in the future (see below), at the moment it should serve as yet another piece of motivation to use *trAIt* alongside domain expert curation rather than as a replacement.

The interactive involvement of domain experts can also improve *trAIt*’s utility. For example, iterating through trait names and descriptions, towards those that capture both the biological meaning and the expected output format (e.g., units for numeric traits, predefined labels for categorical traits), can improve both completeness and accuracy. However, we did not systematically evaluate this practice here.

We found that the consensus model we employed improved accuracy (**Fig. 1**). However, our accuracy metrics depend on what we define as ground truth. It is often impossible to determine ground truth for a “typical” trait value at the species level, or even whether a given value would be widely agreed upon by experts^12^. In this vein, *trAIt* may arguably leverage the “wisdom of the crowd”^13^.

We identify several future directions for future improvement. *trAIt*’s performance is fundamentally constrained by what it can read. In this respect, paywalls remain a major hurdle. But even for open-access papers, *trAIt* works through APIs that can only read text, not figures or supplemental files. To improve literature availability, implementing an authentication system would allow a user with institutional access to retrieve access-restricted papers from subscription-based publishers through PubMed. To improve readability, developing a mechanism to systematically and legally download PDF files of retrieved papers would allow *trAIt* to extract information from figures and supplemental materials.

The implementation of taxon-specific database APIs substantially increased accuracy and completeness (**Fig. 5**). Ideally, *trAIt* would integrate databases and primary literature in parallel to gather trait data. However, a large barrier to this addition is the inconsistent standardization of APIs^14^. Under the assumption that the user has little to no programming experience, it may be challenging to incorporate the querying of additional APIs to *trAIt*’s code. Popular databases like IUCN^10^ and CCKP^11^ would certainly be useful for most taxa, but API formats across databases are inconsistent and must be added individually, limiting how many specific databases can be implemented.

Artificial intelligence is rapidly becoming integrated into many aspects of biological research. As the volume of scientific literature and biological data continues to grow, AI-driven tools offer an increasingly attractive method of extracting and organizing data at scales that would be impractical through manual curation alone. As demonstrated in our case studies, *trAIt* frequently retrieved incorrect or invalid information. While we did not see a correlation between larger datasets and higher inaccuracy rates (**Fig. S1E**), users may be less inclined to error-check thousands of trait-species pairs. At the same time, the increasing reliance on AI in scientific workflows warrants careful consideration. Our conclusion here, also reflected in several ways in *trAIt*’s implementation, is that AI can assist, but not replace, manual curation of species-trait data—striking a balance between time-saving data retrieval, domain expertise and critical evaluation.

## 4 Data Availability

Our code base is publicly available at: https://github.com/harpak-lab/trAIt

## Acknowledgements

We thank Steve Phelps and members of the Harpak Lab for helpful feedback. This work was funded by NIH grant R35GM151108 and a Pew Biomedical Scholarship to A.H.

## Author Contributions

**Srivi Balaji:** Conceptualization, Data Curation, Formal Analysis, Investigation, Methodology, Software, Validation, Visualization, Writing – original draft, Writing – review & editing. **Katherine A. Martinson:** Conceptualization, Data Curation, Formal Analysis, Investigation, Methodology, Resources, Supervision, Validation, Visualization, Writing – review & editing. **Jessica S. Schellenberger:** Data Curation, Writing – review & editing. **Jyotishko Koley:** Data Curation, Writing – review & editing. **Callen M. Inman:** Data Curation, Writing – review & editing. **Hans A. Hofmann:** Data Curation, Writing – review & editing. **Rebecca L. Young:** Conceptualization, Investigation, Methodology, Project Administration, Supervision, Writing – review & editing. **Arbel Harpak:** Conceptualization, Funding Acquisition, Investigation, Methodology, Project Administration, Resources, Supervision, Validation, Writing – review & editing.

## Supplementary Figures

**Figure S1.**
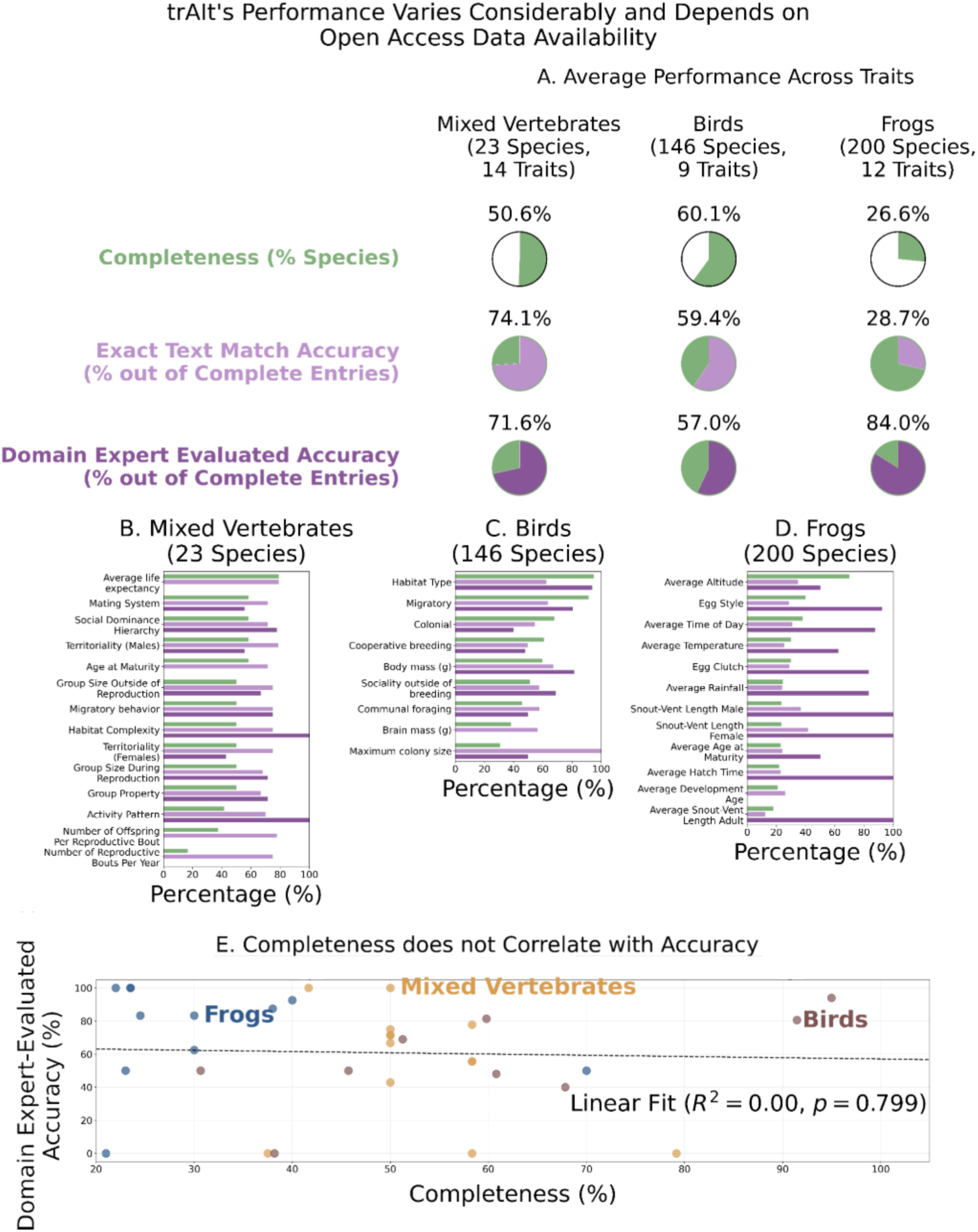
*trAIt* extraction completeness and accuracy metrics across inputs. (A) Summary panel showing average performance across traits for each input (Mixed Vertebrates, Birds, Frogs). For each input, pie charts show completeness (% of species for which *trAIt* returned a non-missing value; green), exact text-match accuracy (% of complete entries matching the ground-truth reference; light purple), and domain expert-evaluated accuracy (% of complete entries evaluated as accurate by a domain expert; dark purple). Note that accuracy metrics are calculated only over complete entries and are therefore not directly comparable to completeness. (B-D) Trait-level breakdown of completeness and accuracy for Mixed Vertebrates (B), Birds (C), and Frogs (D). Birds had the highest level of completeness (60.1%) followed by Mixed Vertebrates (50.6%) and Frogs (26.6%). However, Birds performed moderately using both text-match (59.4%) and domain expert-evaluated accuracy (57.0%), as compared to Mixed Vertebrates, which achieved higher accuracy on both metrics (74.1% on exact text-match and 71.6% evaluated by a domain expert). While Frogs performed poorly when evaluated via exact text match (28.7%), it performed the best of the three inputs when evaluated by a domain expert (84.0%), indicating that values might be inconsistently formatted or be slightly off, evaluated as still correct by a domain expert even if not perfectly matching the reference spreadsheet of values. (E) No significant relationship between completeness and domain expert-evaluated accuracy. Each circle represents a percentage across species for a single trait.

## Notes

https://github.com/harpak-lab/trAIt

### Competing Interest Statement

The authors have declared no competing interest.

